# Comparative single-cell transcriptomics of orthotopic and subcutaneous gastric tumors reveal immune and stromal heterogeneity

**DOI:** 10.1101/2025.10.03.680400

**Authors:** Jinho Jang, Yoojeong Seo, Kyung-Pil Ko, Jie Zhang, Sohee Jun, Jae-Il Park

## Abstract

Preclinical cancer models often use subcutaneous (SC) implantation, which fails to recapitulate the native tumor microenvironment (TME) of orthotopic (ORT) sites. To resolve these differences, we used single-cell RNA sequencing (scRNA-seq) on paired SC and ORT implants of the CKP syngeneic gastric cancer model. Histopathological differences were minimal, but scRNA-seq revealed profound TME divergence. ORT tumors displayed robust stromal activation, coordinated fibroblast and endothelial signaling, and an immune compartment marked by higher T/NK cell activation and IgA-biased B cell plasma programs, reflecting a physiological mucosal environment. In contrast, SC tumors had higher overall T cell infiltration but showed markedly increased CD8+ T cell exhaustion and an enriched oxidative tumor program. Our findings provide critical guidance: SC models are optimal for high-throughput and exhaustion-focused assays, whereas ORT models are indispensable for studying organ-specific immune and stromal biology with translational fidelity.

---

Preclinical mouse models are widely used in cancer research, and the selection of models has significant consequences for study design, interpretation, and downstream translation [1]. Subcutaneous cell line-derived models are commonly used because they are simple, fast, and cost-efficient, and because tumor growth can be quantified easily by serial caliper measurements [2]. However, the subcutaneous site places tumors outside the native organ context, where stromal architecture, vasculature, and immune cues shape tumor behavior and responses to therapy, which limits the use of subcutaneous models for addressing questions related to organ-specific microenvironments [2, 3]. Orthotopic implantation restores the anatomical niche and local microenvironment, and data from such models can more closely reflect clinical disease, at the cost of surgical procedures, specialized monitoring, longer timelines, and potentially greater experimental variability that may require larger cohorts [3].

Anatomical location can condition anti-tumor immunity in vivo [3]. For immunotherapy investigations, a functionally intact immune system is required for meaningful efficacy testing, which generally necessitates syngeneic immunocompetent hosts or carefully designed humanization strategies [4]. Bulk RNA-sequencing of patient-derived xenografts (PDXs) indicates that tumor-intrinsic transcriptional programs are broadly conserved between subcutaneous and orthotopic sites, whereas stromal and other microenvironmental signals diverge [5]. Prior single-cell work has contrasted subcutaneous with intracranial medulloblastoma PDXs, but immune-compartment resolution was limited [6]. Collectively, previous work relied on bulk profiling that obscures cell-type signals [5] and on single-cell analyses with limited immune resolution [6], motivating a comparative single-cell analysis focused on organ-conditioned stromal and immune programs.

Within this context, we used a gastric cancer model as an experimental platform to compare subcutaneous and orthotopic implantation using single-cell RNA sequencing (scRNA-seq). No prior study has paired subcutaneous and orthotopic implants from the same gastric cancer cellular source and profiled them by scRNA-seq with comprehensive immune and stromal annotation.

We utilized a CKP (*Cracd*^-/-^ *Kras*^G12D^ *Trp53*^-/-^) syngeneic model that recapitulates mucinous gastric cancer [7] by subcutaneous (SC) or orthotopic (ORT) implantation. Histopathological examination of CKP-derived tumors revealed mucinous and poorly cohesive components, including signet ring cell carcinoma, in both SC and ORT gastric models. According to WHO/Lauren classification, signet ring cell carcinoma belongs to poorly diffuse type, but no major histopathological differences (**Figure 1**). For these tumors, single-cell RNA-seq libraries were generated and sequenced as previously described [7] (**Figure 2*A***).

**Figure 1.**
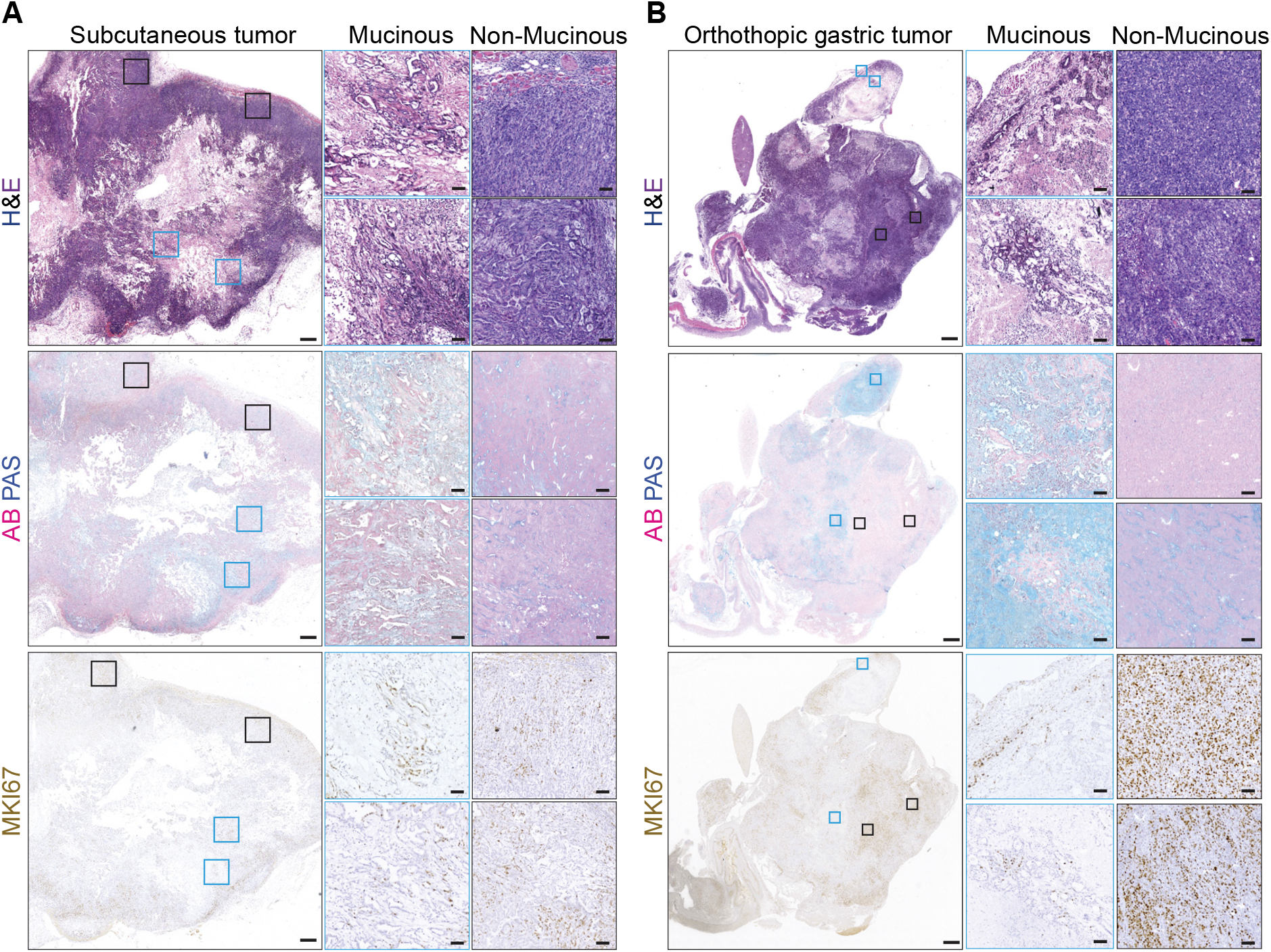
Histological features of subcutaneous and orthotopic gastric tumors. Hematoxylin and eosin (H&E), Alcian blue–periodic acid–Schiff (AB-PAS), and MKI67 staining of *(A)* subcutaneous tumors and *(B)* orthotopic gastric tumors derived from CKP (*Cracd* ^-/-^, *Kras*^G12D^, *Trp53*^-/-^). Blue boxes indicate mucinous regions, and black ones indicate non-mucinous regions. Scale bars: 500 *µ*m; insets, 50 *µ*m. Representative images are shown.

**Figure 2.**
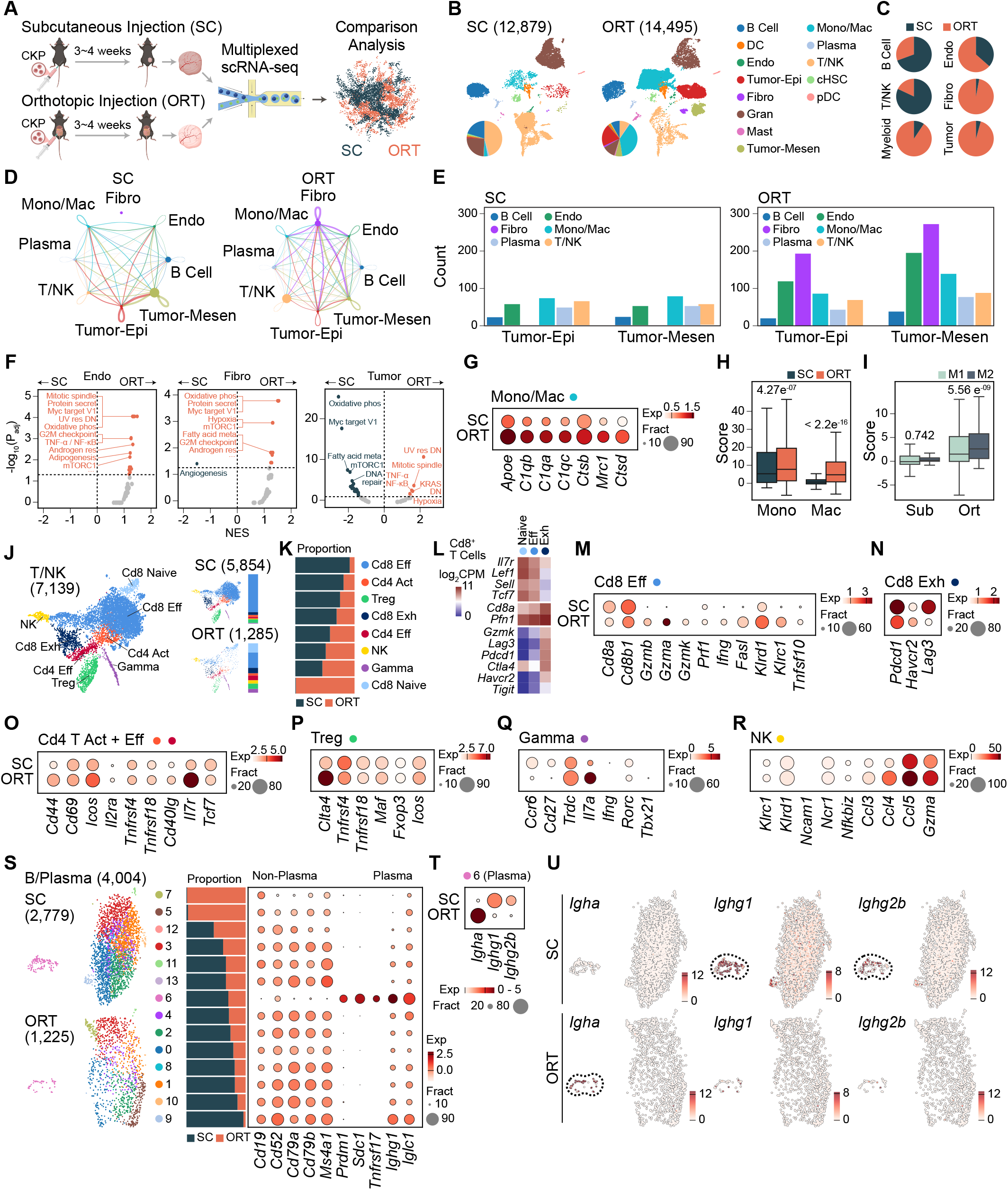
Single-cell profiling reveals site-conditioned TME composition, signaling, and lymphoid programs in CKP tumors. (*A*) Experimental scheme. CKP tumors were implanted subcutaneously (SC) or orthotopically (ORT). Single-cell RNA-seq libraries were generated and jointly analyzed. (*B*) UMAP of all cells with cell-type annotations for SC and ORT; accompanying pie charts show per-sample composition. *(C)* Pie charts showing the fractions of major compartments in SC versus ORT. *(D)* CellChat network maps for SC and ORT among stromal, immune, and tumor compartments. Node size reflects compartment size. Edge width reflects the number of significant ligand-receptor interactions. *(E)* Bar plots of CellChat interaction counts per compartment in SC and ORT. *(F)* Hallmark pathway enrichment comparing ORT versus SC. The x-axis plots the normalized enrichment score (NES), with values above 0 indicating enrichment in ORT and values below 0 indicating enrichment in SC. The y-axis lists the Hallmark gene sets. Separate subpanels display Endo, Fibro, and Tumor. *(G)* Dot plot of macrophage markers in SC and ORT. Dot size indicates the fraction of cells expressing each gene; dot color indicates average expression. *(H)* Box plots of the composite monocyte/macrophage score per cell in SC and ORT, computed from monocyte and macrophage marker sets (center line median; box IQR; whiskers 1.5×IQR; t-test *P*-value shown). *(I)* Box plots of M1-like and M2-like macrophage signature scores per cell in SC and ORT. *(J)* UMAP of T/NK cells with subcluster identities. *(K)* Proportional composition of T/NK subclusters in SC and ORT. *(L)* Heatmap of representative markers used for Cd8^+^ T cell subcluster annotation. *(M-Q)* Dot plots of T cell programs in SC and ORT: (*M)* CD8^+^ effector markers; *(N)* CD8^+^ exhaustion markers; *(O)* CD4^+^ activated/effector markers; *(P)* Treg markers; *(Q)* γδ T cell markers. *(R)* NK cell markers indicating elevated cytotoxic and chemokine programs in ORT (e.g., *Ccl4, Ccl5, Gzma*). *(S)* UMAP of B-lineage cells with subclusters; bar plot of SC and ORT proportions; dot plot showing non-plasma and plasma marker expression. *(T)* Dot plot of plasma isotype programs in SC and ORT: SC upregulates *Ighg1*/*Ighg2b* (IgG), whereas ORT upregulates *Igha* (IgA). *(U)* Feature plots showing *Igha, Ighg1*, and *Ighg2b* expression by condition. ORT plasma shows an IgA bias, and SC plasma shows an IgG bias.

We profiled the tumor microenvironment (TME) in SC and ORT CKP tumors, with emphasis on stromal and immune compartments (**Figure 2*B***). ORT had more tumor cells (epithelial + mesenchymal; 175 SC vs. 3,876 ORT), as well as myeloid cells (602 vs. 5,812) and fibroblasts (6 vs. 201) (**Figure 2*B-C***). In contrast, T/NK (natural killer) cells were more abundant in SC (5,854 SC vs. 1,285 ORT), suggesting higher T cell infiltration in SC (**Figure 2*B-C***). CellChat showed that epithelial/mesenchymal tumor cells engaged in markedly more ligand-receptor interactions with stromal compartments in ORT than in SC (**Figure 2*D-E***). Additionally, fibroblasts and endothelial cells from ORT exhibited broader pathway activation, with more significantly upregulated Hallmark gene sets relative to SC (**Figure 2*F***). By contrast, pathway analysis of epithelial/mesenchymal tumor cells showed enrichment of Hallmark Oxidative Phosphorylation and MYC Targets in SC (**Figure 2*F***). Despite lower tumor cell recovery in SC due to high T cell infiltration, these enrichments indicate a more oxidative and proliferative tumor program of SC independent of overall yield [8]. In parallel, transcription factor activity profiling distinguished SC and ORT tumor cells, revealing distinct regulon activity (**Supplementary Figure 1**). Within the myeloid compartment, ORT exhibited higher global macrophage scores and higher M1/M2-like signature scores, particularly M2-like (**Figure 2*G-I***). Because cell numbers were highly imbalanced, subsequent subclustering was conducted only for T and B cells.

Unsupervised subclustering of T/NK cells resolved a more diverse repertoire in ORT (**Figure 2*J-K***). Notably, CD8^+^ naïve T cells were detected only in ORT (**Figure 2*J-L***). Among T cells, CD8^+^ effector programs showed higher marker expression in ORT, whereas SC exhibited stronger CD8^+^ exhaustion, with increased expression of exhaustion-associated genes within the exhausted CD8^+^ cluster (**Figure 2*M-N***). ORT also showed higher expression of subtype-defining markers in CD4^+^ activated-effector, regulatory T, and γδ T cells, indicating overall higher activation in ORT (**Figure 2*O-Q***). ORT NK cells also showed an increased expression of *Ccl4, Ccl5*, and *Gzma*, consistent with heightened cytotoxicity (**Figure 2*R***). Taken together, these data indicate that ORT better captures activated T/NK biology, whereas the SC preferentially accumulates exhausted CD8^+^ T cells.

The B cell subset was reclustered, resolving discrete transcriptional states and site-dependent isotype programs (**Figure 2*S***). Marker-based annotation indicated that non-plasma B cells were enriched in SC and showed higher expression of non-plasma programs (**Figure 2*S***). Plasma cells from SC showed *Ighg1* and *Ighg2b* upregulation (serum-oriented IgG) [9], whereas *Igha* (mucosal secretory IgA) was increased in ORT plasma cells [10] (**Figure 2*T-U***). These data support site-specific class switching, with an IgA bias in ORT and an IgG bias in SC, consistent with organ-conditioned immunity. Accordingly, ORT provides a physiologic mucosal context for interrogating IgA-oriented B cell programs not recapitulated in SC.

This study utilizes a single syngeneic gastric model with paired SC and ORT implants. For applying to other cancer types, further analysis of other orthotopic models is warranted, along with experimental validation. Uneven cell recovery between sites limited subclustering depth, particularly for the myeloid and fibroblast compartments, which could be overcome by small nuclear RNA sequencing.

Our single-cell profiling of SC and ORT gastric tumors reveals organ-specific TME, with coordinated fibroblast and endothelial activation, activated T/NK states, myeloid polarization, and IgA-biased plasma programs in ORT. SC shows higher T cell abundance with its greater exhaustion. These contrasts provide a crucial guidance for model selection: SC is recommended for high-throughput screening and exhaustion-focused readouts, reserving ORT to explore the complex, organ-specific immune and stromal microenvironment. This study unveils previously underrecognized site-conditioned stromal and immune programs at single-cell resolution.

## Acknowledgments

This work was supported by the Cancer Prevention and Research Institute of Texas (CPRIT) (RP200315 to J.-I.P.) and the National Cancer Institute (CA296049 to J.-I.P.).

## Graphical abstract

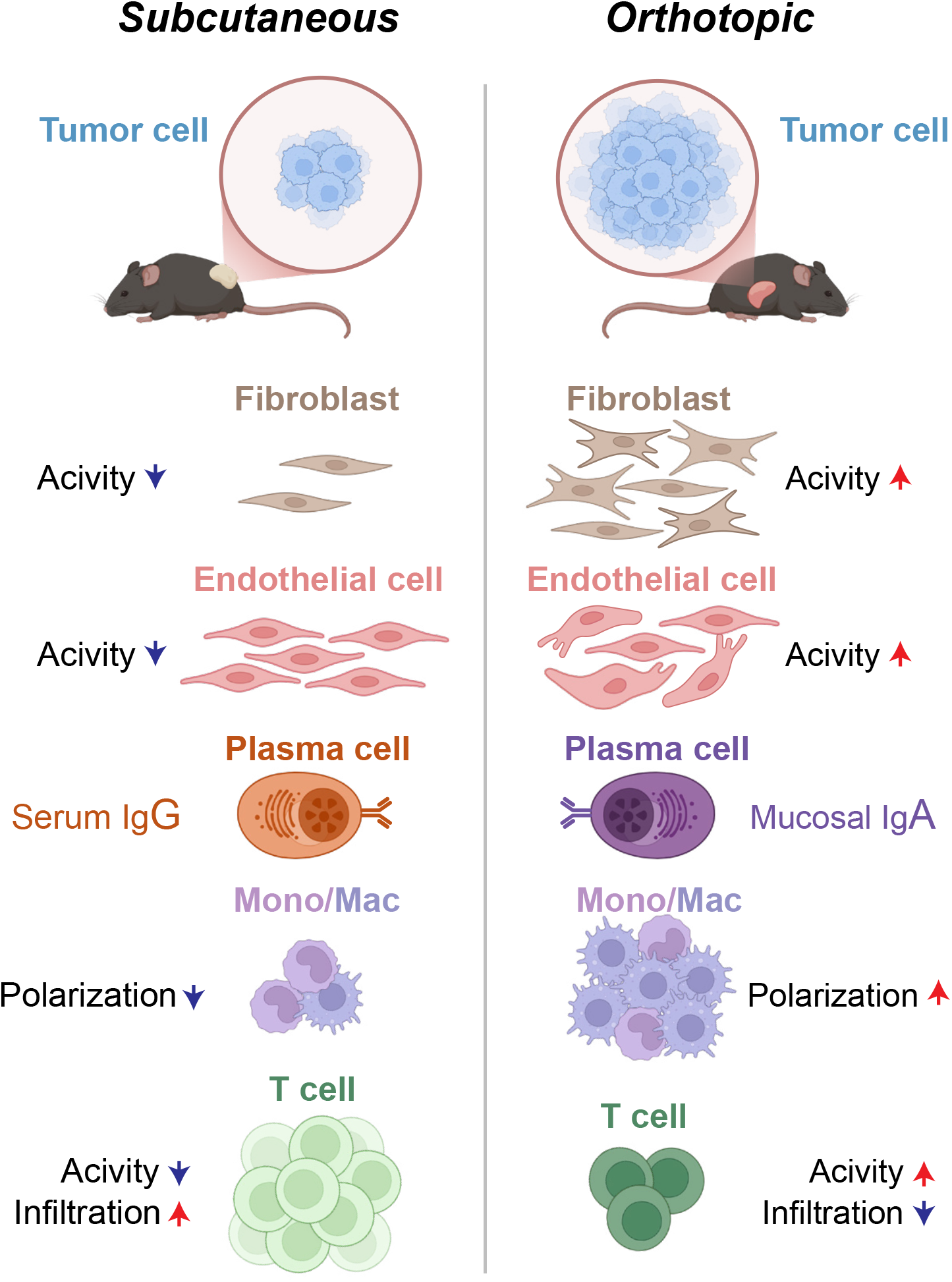

