## Supplementary Materials for "Comparative single-cell transcriptomics of orthotopic and subcutaneous gastric tumors reveal immune and stromal heterogeneity"

### Supplementary Online Methods

*Tumor tissues staining*

All staining was performed as previously described [1]. For immunohistology staining, after blocking with 5% goat serum in phosphate-buffered saline (PBS) for 1 hour at room temperature, sections were incubated with MKI67 antibody (Abcam, ab16667 [1:200]) overnight at 4°C and secondary antibody for 1 hour at room temperature in the dark.

*Subcutaneous and orthotopic injection*

C57BL/6 mice (6-8 weeks) were purchased from the Jackson Laboratory. Mice were randomized and subcutaneously injected with $3 \times 10^6$ cells into both flanks. Mice were maintained in the Division of Laboratory Animal Resources facility at the MD Anderson Cancer Center. Tumor volume was monitored and calculated by measuring with calipers every 2 days (volume = [length x width$^2$] / 2). Tumor burden was determined by measuring all tumor lesions within the lung to account for the complete metastatic load. On days 25-30, mice were euthanized, and tumors were photographed and collected for paraffin embedding followed by immunostaining or single-cell RNA sequencing (scRNA-seq), as previously described [1]. For orthotopic tumor implantation, $1 \times 10^6$ cells were suspended in 50 µl of 50% Matrigel/PBS and injected into the distal gastric wall of C57BL/6 mice (6-8 weeks old). All procedures were conducted in accordance with institutional policies under protocols approved by the Institutional Animal Cancer and Use Committee.

*Processing and integration of scRNA-seq datasets*

scRNA-seq reads were aligned with Cell Ranger (v9.0.1) and matrices were processed in Scanpy [2] (v1.11.1). Identical QC was applied to both subcutaneous and orthotopic datasets: cells with fewer than 100 detected genes and genes detected in fewer than 3 cells were removed; cells were further filtered to mitochondrial content below 20 percent, detected genes below 7,000, and total counts between 0 and 100,000. Doublets were identified with Scrublet [3] (v0.2.3) and excluded, and variation associated with total counts and mitochondrial percentage was regressed out. The datasets were then aligned on 1a6,914 shared genes and concatenated with an implantation-site label (subcutaneous or orthotopic). Data were scaled and subjected to principal component analysis, neighborhood graph construction, and Uniform Manifold Approximation and Projection algorithm (UMAP) embedding [4], followed by Leiden community detection[5] at resolution 2.0. Batch effects were corrected with Harmony [6] using implantation site as the covariate.

*Inference cellular interactions and transcription factor activity*

Cellular interactions were inferred using CellChat [7] (v2.2.0), and transcription-factor regulons were reconstructed with pySCENIC [8] (v.0.12.1) : TF-target modules were first inferred with GRNBoost2 from filtered scRNA-seq, enriched motifs were identified with cisTarget to define regulons, and per-cell regulon activity was scored with AUCell, normalized, and used for clustering and regulon specificity scores.

*Data and code availability*

scRNA-seq data is available via the Gene Expression Omnibus (GEO; accession number: GSE284638; token for reviewers: qdermiqsflmnlep). The code used to reproduce the analyses described in this study can be accessed via Github (https://github.com/jaeilparklab/Ortho_vs_subQ).

*CKP cell lines*

CKP 2D organoid-derived cell lines were maintained in RPMI-1640 + 10% fetal bovine serum (FBS) + 1% penicillin and streptomycin (Pen/Strep) at 37 °C, 5% $CO_2$, routinely confirmed mycoplasma-negative, and used within ≤10 passages for all in vivo experiments [9].

### Supplementary Figure Legends

**Supplementary Figure 1: pySCENIC-derived transcription factor regulon activity in tumor cells from SC and ORT**

Heatmap showing transcription factor regulon activity for tumor epithelial (Tumor-Epi) and tumor mesenchymal (Tumor-Mesen) cells profiled by pySCENIC. Rows are TF regulons. Columns are cells. Values are z-scores of AUCell regulon activity. Clustering was performed on regulons and samples to reveal patterns. Annotation bars indicate implantation site (SC or ORT). The map highlights site-dependent differences in tumor-cell programs, with distinct regulon activity profiles between subcutaneous and orthotopic conditions.

**Supplementary Figure S1**

A

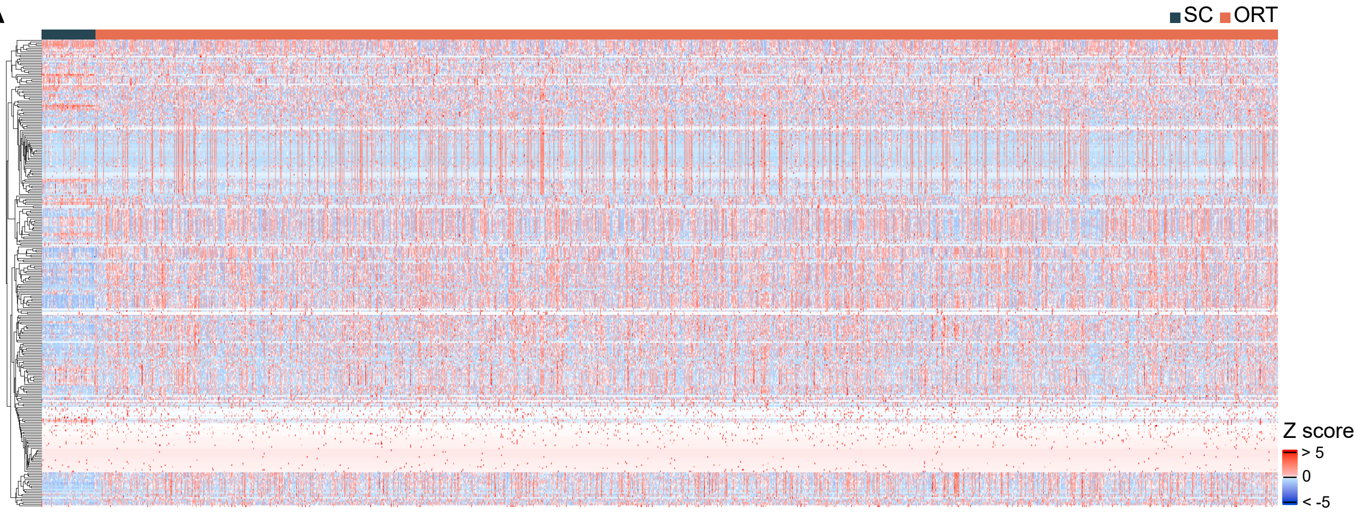
