## Supplementary figures and images for "Comparative single-cell transcriptomics of orthotopic and subcutaneous gastric tumors reveal immune and stromal heterogeneity"

### Supplementary Figure S1

**Supplementary Figure S1**

A

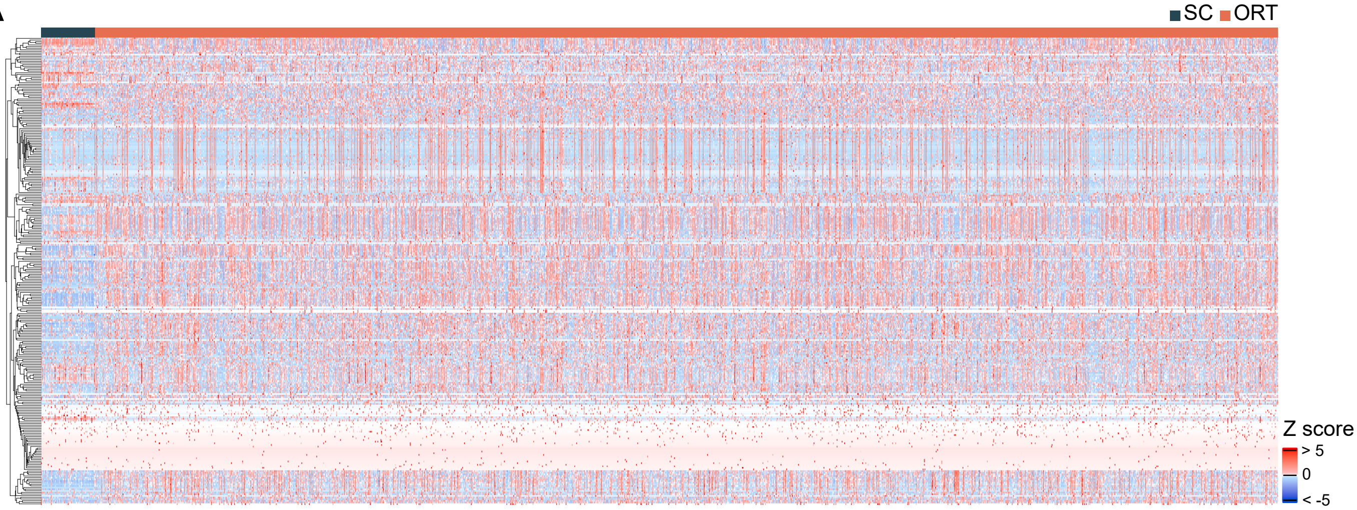
